# Learning Shared Residue Backgrounds and Modification-Specific Offsets for PTM Site Prediction

**DOI:** 10.64898/2026.08.05.743079

**Authors:** Suresh Pokharel, Bishnu Bhusal

**Affiliations:** Rochester Institute of Technology; University of Missouri

**Keywords:** post-translational modifications, protein language models, flow matching, prototype learning, PTM site prediction

## Abstract

Post-translational modifications (PTMs) are chemical changes added to proteins after translation. These changes affect protein function and regulation, and their disruption is linked to disease-associated mechanisms. Because experimentally validating all possible modification sites is impractical, many computational predictors have been developed for PTM site prediction. In this work, we study whether a shared model can represent common residue-background patterns while learning modification-specific background-to-positive offsets. This framing is especially relevant for residues such as lysine (K), which can be acetylated, ubiquitinated, methylated, or sumoylated depending on the surrounding protein context. We propose an anchor-guided rectified flow matching framework for multi-type PTM site prediction from protein language model embeddings. For each PTM–residue pair, the model builds residue-background anchors from PTM-compatible unannotated residues and positive anchors from experimentally annotated modified residues. Given a candidate residue and target modification type, the model compares the residue embedding with these anchor sets and uses a rectified flow module to estimate a modification-conditioned background-to-positive offset. This offset is combined with anchor-based features and used for site scoring. We evaluate the framework on a dbPTM-derived benchmark covering six commonly studied PTMs: phosphorylation, acetylation, ubiquitination, methylation, sumoylation, and N-linked glycosylation. In the shared-model setting, our approach achieves a macro AUPRC of 0.4195, improving over the gated multi-anchor baseline of 0.4154, while independently trained per-modification models achieve 0.4353. These results suggest that multi-type PTM prediction can be modeled within a single shared framework by combining residue-background anchors with modification-conditioned offset features.

## Introduction

Post-translational modifications (PTMs) are covalent changes to proteins that affect protein function and regulation. They are involved in major cellular processes, and their disruption is associated with diverse disease mechanisms Deribe et al. [2010]. Although experimental analysis is the most reliable approach, exhaustive PTM mapping remains challenging because protein sequence space is large and experimental validation is costly and time-consuming Ramazi and Zahiri [2021]. As a result, numerous computational predictors have been developed to identify likely candidate modification sites.

A central difficulty in multi-type PTM prediction is that residue compatibility is shared, but modification identity is specific: the same amino acid type can support multiple PTM types, and different modifications may occur on the same residue site in different biological contexts. For instance, the *ε*-amino group of lysine (K) can be acetylated, methylated, ubiquitinated, or sumoylated Ramazi and Zahiri [2021], Alleyn et al. [2018]. This motivates a shared representation view of multi-type PTM prediction, where common residue-background patterns are learned across PTM types and modification-conditioned offsets capture the patterns that distinguish individual modifications.

The benchmark used in this study alone contains over 1.7 million annotated sites across six PTM types (Table 1), illustrating why training and maintaining a separate model per modification does not scale gracefully as PTM databases continue to grow.

**Table 1:** Summary of PTM types, valid residues, proteins, and annotated positive sites after filtering.

| PTM Type | Residue(s) | Proteins | Annotated Sites |
| --- | --- | --- | --- |
| Phosphorylation | S, T, Y | 187,168 | 1,393,764 |
| Acetylation | K | 34,984 | 121,734 |
| Ubiquitination | K | 31,044 | 154,147 |
| Methylation | K, R | 6,038 | 15,186 |
| Sumoylation | K | 1,681 | 5,773 |
| N-linked glycosylation | N | 6,883 | 26,550 |

Early computational predictors relied on residue windows, physicochemical descriptors, evolutionary profiles, and classical machine-learning models, while later deep-learning methods learned sequence patterns directly from encoded protein fragments Ramazi and Zahiri [2021]. More recent works have used pretrained protein representations, prompt-based learning, multi-task modeling, curated multi-type datasets, and structure-aware residue representations. For example, ProtT5 embeddings have been used as contextual residue features for succinylation prediction Pokharel et al. [2022], PTMGPT2 applies prompt-based fine-tuning across multiple PTM types Shrestha et al. [2024], MTPrompt-PTM uses prompt tuning in a multi-task setting Han et al. [2025], DeepMVP emphasizes large-scale curated PTM data for multi-type prediction Wen et al. [2025], and MeToken combines sequence and structural context Tan et al. [2025]. These studies show substantial progress in PTM prediction, but they primarily improve the input representation, training formulation, or data resource. This work addresses a complementary question: whether a single shared model can predict multiple PTM types by learning common residue-background representations and modification-conditioned background-to-positive offsets.

In this work, we propose an anchor-guided rectified flow matching framework for shared multi-type PTM site modeling in protein language model embedding space using ProtT5 representations. For each PTM–residue pair, the framework constructs residue-background anchors from PTM-compatible candidate residues and positive anchors from experimentally confirmed modified residues. The model uses rectified flow matching to estimate how modification-positive residues deviate from the corresponding residue-background distribution. This learned deviation is combined with anchor-similarity features and used by a shared scoring model to predict whether a candidate site is likely to carry a given PTM.

We evaluate our approach on a dbPTM-derived benchmark across six modification types: phosphorylation (S, T, Y), acetylation (K), ubiquitination (K), methylation (K, R), sumoylation (K), and N-linked glycosylation (N). This setting tests both shared residue-background modeling and PTM-specific behavior, especially for lysine-centered modifications where multiple PTM types share the same residue alphabet. The results suggest that anchor-guided flow matching improves shared anchor-based modeling and supports our hypothesis that multi-type PTM sites can be modeled through a shared residue-background representation together with modification-conditioned background-to-positive offsets.

In summary, this work makes the following contributions:

- We introduce a shared query-based formulation for multi-type PTM site modeling, where PTM-compatible residue backgrounds are separated from modification-specific background-to-positive deviations.
- We propose an anchor-guided rectified flow matching framework that constructs background and modification-positive anchors and uses learned offset features for PTM-aware site scoring within a single shared model.
- We provide a controlled empirical study across six PTM types, showing when shared anchor-flow modeling is competitive with PTM-specific training and analyzing the effect of anchor-guided flow matching relative to anchor-based modeling alone.

## Related Work

### PTM site prediction

Computational PTM site prediction has evolved from motif-based and feature-engineered methods to classical machine learning and deep neural networks Ramazi and Zahiri [2021], Zhang et al. [2026]. Earlier predictors commonly used local sequence windows, physicochemical descriptors, evolutionary profiles, and modification-specific models. More recent deep learning methods learn sequence representations directly, but the task remains challenging because modified sites are sparse, valid residue alphabets differ across modification types, and positive annotations are uneven across PTM categories Audagnotto and Dal Peraro [2017].

### Protein language models for PTM prediction

Protein language models are trained on large corpora of unlabeled protein sequences and/or structures and provide contextual protein representations and have become widely used in protein function prediction, including PTM prediction. In PTM site prediction, embeddings from models such as ESM, ProtTrans, and SaProt are commonly used as input features for downstream predictors. For example, LMSuccSite combined ProtT5 (a ProtTrans model) embeddings with supervised word embeddings to predict succinylation sites Pokharel et al. [2022].

Recent work has explored prompt-based, multi-task, and multi-type formulations that go beyond using frozen embeddings as static input features. PTMGPT2 fine-tunes a GPT-2-based protein language model with PTM-specific prompts and uses attention visualization for interpretability, evaluated across 19 PTM types Shrestha et al. [2024]. MTPrompt-PTM instead applies prompt tuning to a structure-aware protein language model within a multi-task framework, combining shared feature-extraction layers with task-specific heads and a knowledge-distillation strategy across 13 PTM types Han et al. [2025]. DeepMVP takes a data-centric approach, training on a curated resource of nearly 400,000 PTM sites obtained by reprocessing hundreds of mass-spectrometry datasets, for the same six PTM types studied in this work Wen et al. [2025]. MeToken instead tokenizes the sequence and structural micro-environment of each residue into discrete, PTM-aware tokens, and specifically targets the long-tailed distribution of PTM types through uniform sub-codebooks so that rare modifications remain well represented Tan et al. [2025].

These methods primarily improve performance by fine-tuning or prompt-tuning the underlying language model, incorporating structural context, or scaling up curated training data. Our approach instead keeps the pretrained ProtT5 backbone frozen and addresses a complementary question: whether a single shared model can use background and modification-positive anchor regions, together with a learned flow-based offset, to represent residue-level PTM differences without modifying the language model itself or requiring structural input. This anchor-and-offset formulation is lighter-weight than fine-tuning a language model separately for each PTM type, though, unlike MeToken, it does not include an explicit mechanism for the long-tailed imbalance across PTM types, which Table 1 shows is substantial in our dataset as well.

### Anchor-based representation learning

Prototype-based learning represents classes using prototype embeddings, often computed from support examples, and predicts query labels using distances or similarities to the prototypes Snell et al. [2017]. Our approach follows this reference-based approach, but adapts it to residue-level PTM modeling. Rather than representing each modification only through the downstream prediction head, we use anchors as compact, prototype-like summaries of PTM-compatible background residues and experimentally observed modification-positive residues. These anchors provide reference points for estimating how modified residues deviate from the background residues.

### Flow matching and rectified flow

Flow matching and rectified flow learn vector fields that map samples between distributions and have been widely studied in generative modeling Lipman et al. [2023], Liu et al. [2023]. Recent work has also discussed flow matching as a framework for learning mappings between high-dimensional biological data distributions Morehead et al. [2026]. We use this idea in a discriminative setting. Rather than generating new protein embeddings, the trained flow module estimates modification-conditioned background-to-positive offset features. These features describe how a candidate residue deviates from its residue-background anchors relative to modification-positive anchors, and are used for downstream PTM site prediction.

## Methodology

### Dataset and Candidate Sites

We collected experimentally annotated PTM sites from dbPTM, a curated database of protein post-translational modifications Chung et al. [2025]. We focused on six common PTM types with well-defined residue specificity: general phosphorylation on serine (S), threonine (T), and tyrosine (Y); acetylation, ubiquitination, and sumoylation on lysine (K); methylation on lysine (K) and arginine (R); and N-linked glycosylation on asparagine (N). Proteins were retained only when a full-length ProtT5 embedding was available and the UniProt identifier matched exactly. A summary of the filtered dataset is shown in Table 1.

Candidate sites were defined using PTM-specific residue compatibility. For example, phosphorylation candidates were all serine, threonine, and tyrosine residues, whereas acetylation, ubiquitination, and sumoylation candidates were lysine residues; methylation candidates were lysine and arginine residues; and N-linked glycosylation candidates were asparagine residues. Experimentally annotated sites were labeled as positives. Compatible residues without dbPTM annotations were treated as candidate negatives for supervised training and evaluation, recognizing that unannotated residues may include undiscovered modification sites.

To avoid protein-level information leakage, we split the data by protein rather than by residue. All unique proteins appearing in any of the six PTM datasets were assigned to training, validation, and test sets using a global 80:10:10 split with random seed 42. Thus, all candidate and annotated sites from the same protein appear in only one split, even when the protein contains multiple PTM types.

### Problem Formulation

We formulate multi-type PTM site prediction as a residue-level binary query problem. Given a protein *P*, a residue position *i*, its residue identity *r*_*i*_, the corresponding residue’s representation *x*_*i*_, and a PTM type *k*, the goal is to estimate

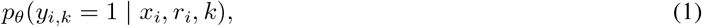

where *y*_*i,k*_ = 1 indicates that residue *i* is experimentally annotated with PTM type *k*, and *y*_*i,k*_ = 0 indicates a PTM-compatible but unannotated candidate residue, which is commonly considered a negative site.

Candidate residues are restricted to the valid residue alphabet for each PTM type. For example, phosphorylation is queried only on serine (S), threonine (T), and tyrosine (Y) residues; acetylation, ubiquitination, and sumoylation are queried on lysine (K) residues; methylation is queried on lysine (K) and arginine (R) residues; and N-linked glycosylation is queried on asparagine (N) residues. Thus, the model scores only PTM-compatible site–PTM queries rather than all possible residue–PTM combinations.

Each valid site–PTM query (*i, k*) is modeled using a shared residue-background anchor 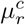 and a PTM-specific background-to-positive deviation 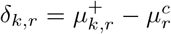. As illustrated in Figure 1, this formulation allows our framework to share residue-background information across PTM types while still producing modification-specific site scores.

**Figure 1.**
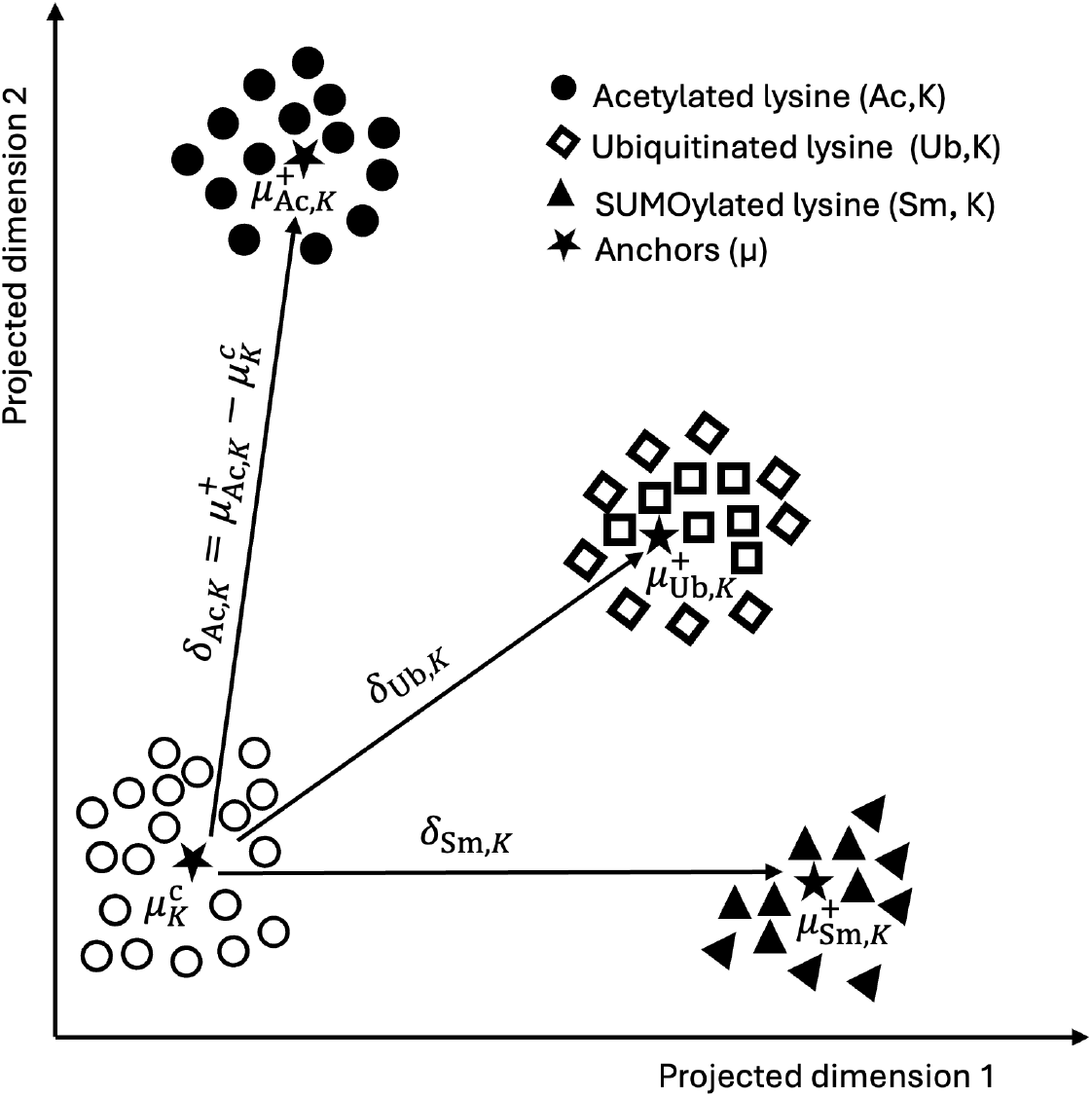
Conceptual illustration of shared residue-background anchors and PTM-specific offsets for lysine-centered modifications. A common background anchor centroid 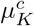 summarizes PTM-compatible lysine candidates, while*k,K*each positive anchor centroid 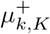 summarizes experimentally annotated sites for PTM type *k*. The offsets 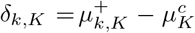 represent modification-conditioned background-to-positive deviations. The plot is schematic; for visual*K*simplicity, each anchor set is shown as a single representative centroid, and the plot is not an actual embedding projection.

### Residue Embeddings

Each protein sequence is represented using contextual residue embeddings from the encoder part of a pretrained ProtT5 protein language model. It is a transformer-based protein language model trained on large-scale protein sequences using self-supervised objectives. For residue position *i*, the corresponding embedding is denoted as

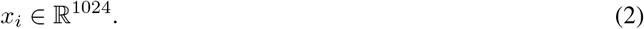

Because embeddings are generated from the full-length sequence, each residue representation reflects broader protein context rather than only a fixed local window.

### Residue-Background and PTM-Positive Anchors

For each residue type *r*, we collect unannotated PTM-compatible training residues of type *r* into a residue-background pool and summarize it with *B* background anchors,

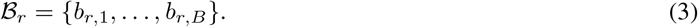

These background anchors are computed independently of PTM type *k*, allowing _*r*_ to serve as a shared residue-background reference. For each PTM–residue pair (*k, r*), we collect experimentally annotated positive training sites and summarize them with *M* positive anchors,

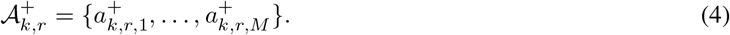

Anchors are computed as cluster centroids obtained by mini-batch *k*-means clustering, fit separately for each group: residue type *r* for background anchors and PTM–residue pair (*k, r*) for positive anchors. Clustering is performed after residue-wise standardization of ProtT5 embeddings,

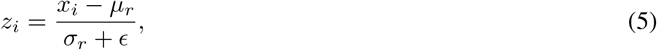

where *µ*_*r*_ and *σ*_*r*_ are estimated from the training background pool of residue type *r*. All anchors and normalization statistics are estimated from the training split only, with no validation or test information. In the main configuration, we use *M* = 16 positive anchors and *B* = 32 background anchors per group.

### Anchor-Guided Rectified Flow Matching

Figure 2 summarizes the overall architecture described in this and the following subsection. As introduced above, anchors define residue-background and PTM-positive reference regions in the standardized embedding space. Now, we use these anchors to construct training pairs for a rectified flow module that learns modification-conditioned background-to-positive deviations. The flow field is a single shared conditional model,

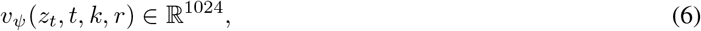

where *z*_*t*_ is a standardized residue embedding, *t* ϵ [0, 1] is the interpolation time, and *k* and *r* denote the PTM type and residue type. PTM and residue identities are represented using learned embeddings and concatenated with *z*_*t*_ and *t* as input to the flow network.

**Figure 2.**
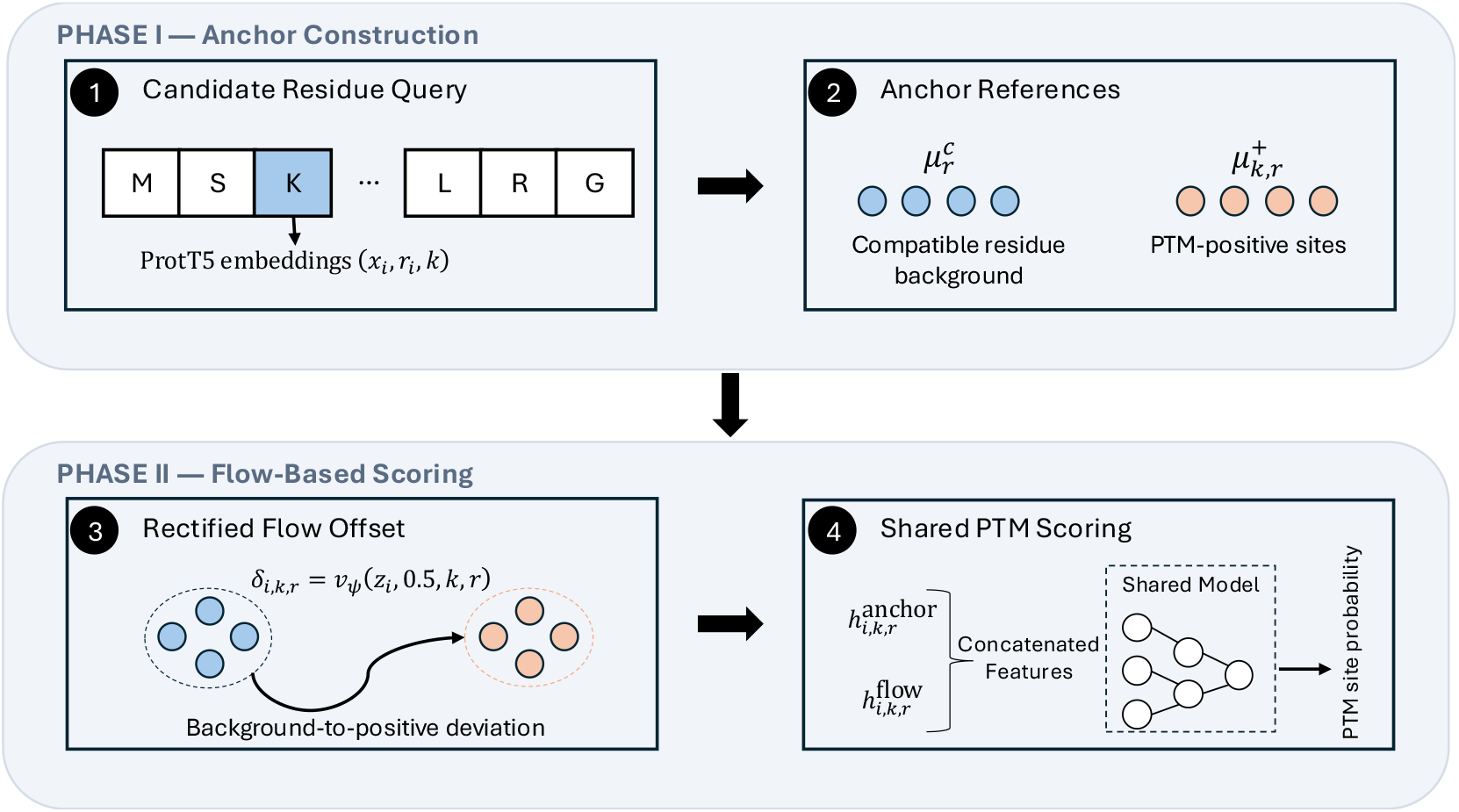
Overview of the proposed framework. For a candidate residue query (*x*_*i*_, *r*_*i*_, *k*), ProtT5 embeddings are used to obtain a standardized residue representation. The model then compares the candidate with residue-background and PTM-positive anchor references to estimate a modification-conditioned background-to-positive deviation *δ*_*i,k,r*_ = *v*_*ψ*_(*z*_*i*_, 0.5, *k, r*) which is later combined with anchor-contrast and flow-derived features in a shared scoring model to predict the site–PTM probability 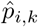.

For each PTM–residue pair (*k, r*), flow pairs are constructed from training embeddings using the anchors. Background embeddings are assigned to their nearest background anchor in B_*r*_, and positive embeddings are assigned to their nearest positive anchor in 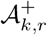. Each positive anchor is then matched to its nearest background anchor by Euclidean distance in standardized space. For each matched anchor pair, *z*_0_ is sampled from the background embeddings assigned to the matched background anchor, and *z*_1_ is sampled from the positive embeddings assigned to the corresponding positive anchor. Thus, the flow is trained on real standardized embeddings, while anchors guide the pairing.

The flow field is trained with the rectified flow matching objective. Given a standardized background embedding *z*_0_ and a matched standardized PTM-positive embedding *z*_1_, we sample

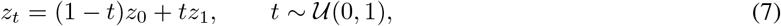

and optimize

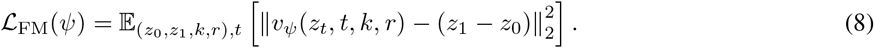

Shared Residue Backgrounds and Modification-Specific Offsets for PTM Site Prediction

This trains the shared conditional flow field to estimate the background-to-positive offset for each PTM–residue context.

At inference, the trained flow field is evaluated once at a fixed value *t*_eval_ = 0.5 using the standardized candidate embedding *z*_*i*_:

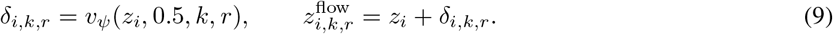

The predicted deviation *δ*_*i,k,r*_ and transported embedding 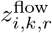 are used as flow-derived features. We also compute scalar summaries including the deviation norm, the cosine similarity between *z*_*i*_ and *δ*_*i,k,r*_, the nearest positive-anchor distance before transport, the nearest positive-anchor distance after transport, and their difference. Each of the two 1024-dimensional flow vectors is projected independently to a 128-dimensional representation, so together with the five scalar summaries the flow module contributes 261 dimensions to the final feature vector. These features are concatenated with the anchor-contrast features and passed to the final PTM scoring model.

### PTM Scoring

After the rectified flow module is trained, it is used as a frozen feature extractor for downstream PTM site scoring. For each valid site–PTM query (*i, k*), we combine two sources of information: anchor-contrast features derived from the background and PTM-positive anchors, and flow-derived features from the predicted background-to-positive deviation.

The flow-derived features include the predicted deviation *δ*_*i,k,r*_, the transported embedding 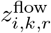, the deviation norm, the cosine similarity between *z*_*i*_ and *δ*_*i,k,r*_, the nearest positive-anchor distance before and after transport, and their difference. The high-dimensional flow vectors are first projected to a lower-dimensional representation and then concatenated with the anchor-contrast features.

The combined representation is passed through a shared gated scoring model that produces a binary logit for the queried site–PTM pair:

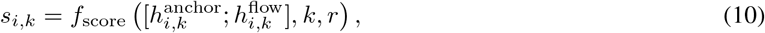

where 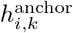 denotes the anchor-contrast representation and 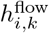 denotes the flow-derived representation. Concretely, *f*_score_ is a shared two-layer feed-forward network (hidden sizes 512 and 128, with ReLU activations and dropout) whose penultimate representation is multiplicatively modulated by a sigmoid gate computed from a learned 32-dimensional PTM embedding, before a final linear layer produces the site–PTM logit. This gating mechanism lets the shared backbone learn cross-PTM structure while still allowing modification-specific rescaling of the shared features. The predicted PTM probability is

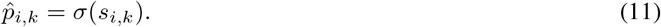

### Experimental Setup

#### Compared Methods

We compare three models that separate the effects of PTM-specific training, shared anchor-based modeling, and anchor-guided flow features.

#### Per-PTM baseline

This baseline trains an independent MLP for each PTM type using only the standardized ProtT5 residue embedding. It provides a strong PTM-specific reference without anchors, flow features, or parameter sharing across PTMs.

#### Gated multi-anchor model

This shared model uses the same residue-background anchors B_*r*_ and PTM-positive anchors 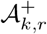 as our full framework, but scores candidates using only anchor-contrast features. It therefore tests the value of anchor-based shared modeling without rectified-flow offsets.

#### Anchor-guided rectified flow matching (full model)

Our full framework adds anchor-guided rectified-flow features to the shared anchor representation. The flow field is trained once across all PTM–residue pairs and then used as a frozen feature extractor for final PTM site scoring.

#### Implementation Details

Each protein sequence of length *L* is encoded using ProtT5, producing a residue-level embedding matrix *X*∈ R^*L×*1024^, as described in the Residue Embeddings section above. The standardization statistics are computed from the training split only and applied to validation and test sets. The main configuration uses *M* = 16 PTM-positive anchors and *B* = 32 residue-background anchors, computed by mini-batch *k*-means in standardized embedding space. Flow pairs are constructed using nearest-neighbor anchor matching, and the trained flow field is evaluated at *t*_eval_ = 0.5.

All models are trained with binary cross-entropy loss using AdamW. During training, negative candidates are under-sampled to a 1:1 ratio with positives to address the class imbalance; validation and test evaluation use the full natural class distribution without undersampling. Models are trained for up to 20 epochs with early stopping (patience of 5 epochs, monitored on validation macro AUPRC), a batch size of 1024, a learning rate of 1 × 10^*−*3^, and a weight decay of 1 × 10^*−*4^.

### Metrics

We use macro AUPRC as the primary metric because PTM site prediction is highly imbalanced and candidate-site counts vary substantially across PTM types. AUPRC is more informative than accuracy in this setting because positive PTM sites are sparse relative to the number of compatible candidate residues. Macro AUPRC is computed by first calculating AUPRC separately for each PTM type and then averaging across PTMs, giving each modification equal weight regardless of dataset size.

We also report pooled AUPRC, which evaluates all site–PTM predictions together, and macro AUROC as an additional ranking-based metric. To assess threshold-dependent prediction quality, we report macro F1 and macro MCC. For each of these two metrics, the classification threshold is selected by sweeping 200 candidate values in [0.01, 0.99] on the validation set and choosing the value that maximizes the corresponding metric; test labels are never used for threshold selection. Per-PTM AUPRC is included to show whether aggregate trends are consistent across modification types or driven by a small number of PTMs.

## Results

### Overall Performance

Table 2 reports the held-out test performance. The independent per-PTM baseline remains the strongest overall reference, with a macro AUPRC of 0.4353 ± 0.0021. Among shared models, our full framework achieves the highest mean macro AUPRC, 0.4195 ± 0.0040, compared with 0.4154 ± 0.0024 for the gated multi-anchor model.

**Table 2:** Held-out test performance for the main compared methods. Values are reported as mean ± standard deviation over five random seeds.

| Method | Macro AUPRC | Pooled AUPRC | Macro AUROC | Macro F1 | Macro MCC |
| --- | --- | --- | --- | --- | --- |
| Per-PTM baseline | **0.4353 ± 0.0021** | 0.3346 ± 0.0075 | **0.8489 ± 0.0010** | **0.4145 ± 0.0094** | **0.3887 ± 0.0071** |
| Gated multi-anchor | 0.4154 ± 0.0024 | **0.3361 ± 0.0050** | 0.8389 ± 0.0012 | 0.3889 ± 0.0174 | 0.3650 ± 0.0129 |
| Anchor-guided flow matching (ours) | 0.4195 ± 0.0040 | 0.3354 ± 0.0038 | 0.8388 ± 0.0018 | 0.4062 ± 0.0110 | 0.3769 ± 0.0079 |

### Effect of Anchor-Guided Flow Matching

To isolate the effect of rectified flow matching, we compare our full framework with the gated multi-anchor model, which uses the same residue-background and PTM-positive anchors but removes the flow field. Anchor-guided flow matching improves macro AUPRC from 0.4154 to 0.4195, a gain of 0.0041. It also improves macro F1 from 0.3889 to 0.4062 and macro MCC from 0.3650 to 0.3769.

The pooled AUPRC is nearly unchanged, decreasing slightly from 0.3361 to 0.3354, and macro AUROC remains essentially flat. This suggests that the flow-derived features do not improve all ranking metrics uniformly, and instead mainly benefit macro-averaged and threshold-dependent metrics such as F1 and MCC.

### Modification-Specific Behavior

Table 3 shows that the effect of flow matching varies across PTM types. Relative to the gated multi-anchor model, anchor-guided flow matching improves four of the six categories, with the largest gains for methylation and sumoylation. Phosphorylation and ubiquitination show small decreases. Compared with the independent per-PTM baseline, our full framework is higher only for phosphorylation and remains lower for the other five PTMs, consistent with the overall result that independent PTM-specific training remains the strongest reference.

**Table 3:** Per-PTM AUPRC comparison across the per-PTM baseline, gated multi-anchor model, and anchor-guided flow matching. Values are means across five random seeds. Bold indicates the best result for each PTM type.

| PTM Type | Per-PTM | Gated | Flow (ours) |
| --- | --- | --- | --- |
| Phosphorylation | 0.3160 | **0.3217** | 0.3204 |
| Acetylation | **0.3249** | 0.3149 | 0.3166 |
| Ubiquitination | **0.3979** | 0.3890 | 0.3874 |
| Methylation | **0.3092** | 0.2706 | 0.2864 |
| Sumoylation | **0.3731** | 0.3261 | 0.3327 |
| N-linked glycosylation | **0.8908** | 0.8703 | 0.8737 |

### Stability Across Seeds

Anchor-guided flow matching improves over the gated multi-anchor model in four of the five seed runs and has the higher mean macro AUPRC. The gated multi-anchor model has lower variance, but its mean performance is lower. This supports using anchor-guided flow matching as the main shared model while keeping the per-PTM baseline as the strongest overall reference.

## Discussion

These results indicate that combining shared residue-background anchors with anchor-guided flow matching is a viable strategy for multi-type PTM site prediction, and that the flow module’s benefit is concentrated in macro-level and threshold-sensitive metrics rather than pooled ranking performance. This is consistent with the relative size of the flow contribution: the projected flow features add only 261 dimensions to a base anchor-contrast feature vector of 8,198 dimensions, so the flow module acts as a comparatively small refinement of the shared anchor representation rather than a dominant input to the scoring model.

The independent per-PTM baseline remains the strongest overall reference, which is expected because each model is optimized for one modification type without sharing parameters across heterogeneous PTM categories. However, our full framework reaches competitive performance while using a single shared framework across all six PTM types.

This comparison is important because the per-PTM baseline trains separate models for each modification, whereas our approach shares the residue-background representation and scoring framework. The relatively small gap to per-PTM training suggests that much of the PTM-relevant signal can be captured through shared residue-background anchors together with modification-conditioned offset features. This is especially relevant for PTMs that share candidate residues, such as lysine-centered modifications, but differ in the embedding-space patterns associated with PTM-positive sites.

The method provides a controlled way to study how PTM-positive sites deviate from compatible residue backgrounds in protein language model embedding space. At the same time, the remaining gap to per-PTM training suggests that future shared models may need stronger mechanisms for PTM-specific adaptation while preserving the stability of a unified framework.

## Limitations

This study has several limitations. First, our framework is evaluated on six commonly studied PTM types for which candidate residues can be defined directly from amino acid identity. Although these PTMs are among the most studied modifications, they represent only a small subset of the reported PTM landscape, and extending the framework to comprehensive PTMs remains future work.

Second, the model relies on sequence-derived ProtT5 embeddings and does not explicitly incorporate structural or biological context such as solvent accessibility, disorder, or enzyme specificity. Structure-aware protein language models or additional biological features may provide complementary information. Furthermore, the evaluation is based on one dbPTM-derived benchmark. We use protein-level splits to reduce information leakage, but broader validation on independent datasets and time-split benchmarks is needed to better assess the generalization of the approach.

Finally, compatible residues that are not annotated as positive in the same sequence are considered negatives, although some may correspond to undiscovered PTM sites. We also focus on controlled internal baselines rather than a comprehensive comparison with public PTM prediction tools, since existing tools often use different datasets, PTM definitions, filtering rules, and evaluation protocols. In addition, the F1 and MCC thresholds are selected independently on the validation set for each metric and each trained model, which is standard practice but introduces some threshold-selection flexibility relative to a single fixed operating point; AUPRC and AUROC, which are threshold-independent, are therefore emphasized as the primary metrics in this study.

## Conclusion

We introduced an anchor-guided rectified flow matching framework for shared multi-type PTM site modeling, which separates PTM-compatible residue backgrounds from modification-specific background-to-positive deviations for PTM-aware site scoring within a single shared model. Across six PTM types, our approach achieves competitive performance relative to independently trained per-PTM models while improving over the shared gated anchor baseline, supporting the premise that multi-type PTM sites can be modeled through a shared residue-background representation together with modification-conditioned offset features.

### Social Impact

PTM dysregulation is associated with major diseases, including cancer and neurodegenerative disorders, and faster identification of candidate PTM sites can support biological discovery. Our framework is designed as a computational prioritization tool for experimental follow-up, not as a replacement for laboratory validation. By ranking candidate sites across multiple PTM types within a single shared model, it may help researchers focus validation efforts on higher-priority residues, potentially reducing experimental time, reagent use, labor, and cost when used together with expert judgment.

The model should be interpreted with caution because reliability is not uniform across PTM types. As shown in Table 1, annotation counts vary substantially across modifications, and Table 3 shows corresponding differences in per-PTM performance. Predictions for underrepresented or harder-to-model PTMs should therefore be treated as candidate rankings rather than confirmed annotations.

The shared design also has practical benefits. Instead of training and maintaining separate models for each PTM type, our framework uses a unified anchor-flow approach that can be extended to additional PTMs when sufficient annotations are available. In future work, we plan to provide a web server so researchers can submit protein sequences and obtain PTM site predictions to guide downstream experiments. Such outputs will be presented as prioritization signals to reduce the risk of unvalidated predictions being treated as experimentally confirmed findings.

## Code Availability

An implementation of the framework described in this paper, including preprocessing, training, and evaluation scripts, is available at: https://github.com/suresh-pokharel/PTM-AnchorFM.

## Acknowledgements

The author thanks the RIT Research Computing for providing the research environment and computational resources that supported this work.

