## Supplementary Information for "Learning Shared Residue Backgrounds and Modification-Specific Offsets for PTM Site Prediction"

[2]University of Missouri

#### A Method Details

##### A.1 Anchor-Conditioned Site Representation

For each valid site–PTM query $(i, k)$, PTM-AnchorFM computes candidate-specific summaries from the residue-background anchors $\mathcal{B}_r$ and PTM-positive anchors $\mathcal{A}_{k,r}^{+}$. These summaries compare the candidate residue with the compatible residue background and the PTM-positive region.

The anchor-conditioned representation combines the standardized residue embedding $z_i$, the positive-anchor summary $c_{i,k,r}^{+}$, the background-anchor summary $c_{i,r}^{0}$, and interaction features such as differences and element-wise products:

$$h_{i,k,r}^{\text{anchor}} = \phi_{\text{anchor}} \left( z_i, c_{i,k,r}^{+}, c_{i,r}^{0} \right). \tag{1}$$

This representation is used by the gated multi-anchor baseline and is concatenated with flow-derived features in PTM-AnchorFM.

##### A.2 Gated Multi-Anchor Baseline

The gated multi-anchor model is the shared anchor-based baseline used in the main paper. It uses the same residue-background and PTM-positive anchors as PTM-AnchorFM, but does not include the rectified flow module. The anchor-conditioned representation is passed through a shared multilayer perceptron, and a learned PTM-conditioned gate modulates the hidden representation before binary site–PTM scoring:

$$s_{i,k}^{\text{anchor}} = f_\theta \left( g_k \odot h_{i,k,r}^{\text{anchor}} \right), \tag{2}$$

where $g_k$ is a learned PTM-specific gate and $s_{i,k}^{\text{anchor}}$ is the logit for the queried site–PTM pair. This baseline isolates anchor-based shared modeling before adding rectified flow features.

| Seed | Gated Anchor | PTM-AnchorFM | PTM-head + RFM |
|---|---|---|---|
| 1 | 0.4134 | 0.4260 | 0.4044 |
| 7 | 0.4170 | 0.4202 | 0.4118 |
| 21 | 0.4188 | 0.4156 | 0.4201 |
| 42 | 0.4149 | 0.4173 | 0.4238 |
| 123 | 0.4131 | 0.4186 | 0.4219 |

Table 1: Seed-level macro AUPRC for shared models and the PTM-specific-head RFM extension.

| Method | Macro AUPRC | Pooled AUPRC | Macro AUROC | Macro F1 | Macro MCC |
|---|---|---|---|---|---|
| PTM-specific head + RFM | $0.4164 \pm 0.0081$ | $0.3357 \pm 0.0047$ | $0.8397 \pm 0.0017$ | $0.3860 \pm 0.0154$ | $0.3632 \pm 0.0125$ |
| PTM-specific head, no RFM | 0.4146 | 0.3319 | 0.8380 | 0.4039 | 0.3753 |
| Local-density RFM | 0.4172 | 0.3290 | 0.8373 | 0.4029 | 0.3746 |

Table 2: Ablation results. Five-seed results are reported as mean $\pm$ standard deviation; single-seed ablations are reported as point estimates.

##### A.3 Flow Pair Construction

For each PTM–residue pair $(k, r)$, background embeddings are assigned to their nearest background anchor in $\mathcal{B}_r$, and positive embeddings are assigned to their nearest positive anchor in $\mathcal{A}^+_{k,r}$. Each positive anchor is then matched to its nearest background anchor in standardized embedding space. For each matched anchor pair, flow training pairs $(z_0, z_1)$ are sampled from the background and positive embeddings assigned to the corresponding clusters. Thus, the rectified flow model is trained on real standardized residue embeddings, while anchors determine the pairing.

#### B Experimental Results

##### B.1 Seed-Level Results

Table 1 reports seed-level macro AUPRC for the main shared models and the PTM-specific-head RFM extension.

##### B.2 Ablation Results

Table 2 reports ablations omitted from the main comparison table.

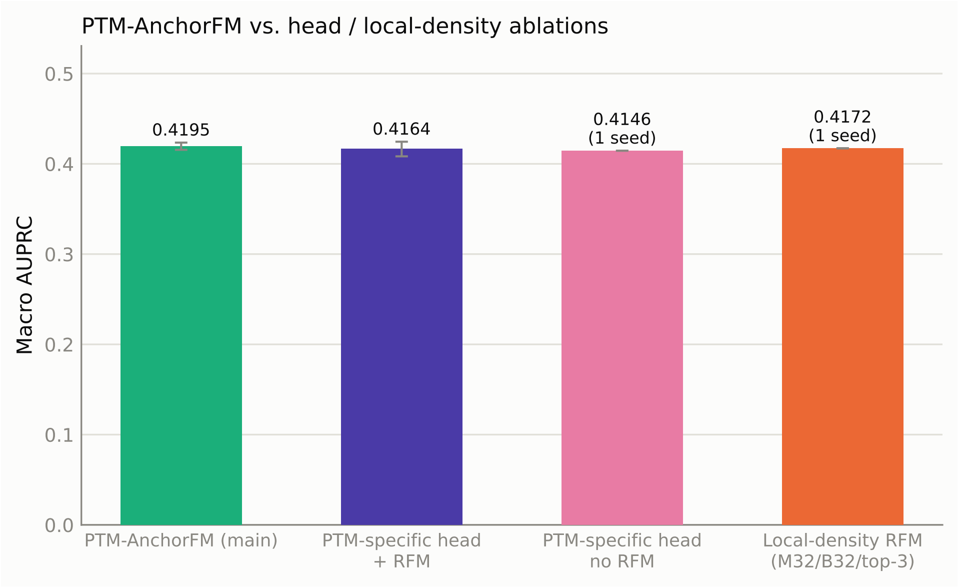


Figure 1: Macro AUPRC comparison between PTM-AnchorFM and head/local-density ablations.

##### B.3 Per-PTM Performance

Figure 2 reports per-PTM AUPRC across models with seed-level variation.

#### C Embedding-Space Visualizations

##### C.1 Lysine-Centered PTM Projections

Figure 3 shows two-dimensional PCA projections for lysine-centered PTMs. Each panel includes standardized ProtT5 residue embeddings, learned background anchors, PTM-positive anchors, and a subsample of flow vectors. These projections are qualitative visualizations; the axes are not directly comparable across panels.

##### C.2 Real Flow Vectors for Methylation

Figure 4 shows the frozen rectified-flow field for methylation on lysine. Arrows are evaluated at $t_{\text{eval}} = 0.5$ and projected onto the top two PCA components.

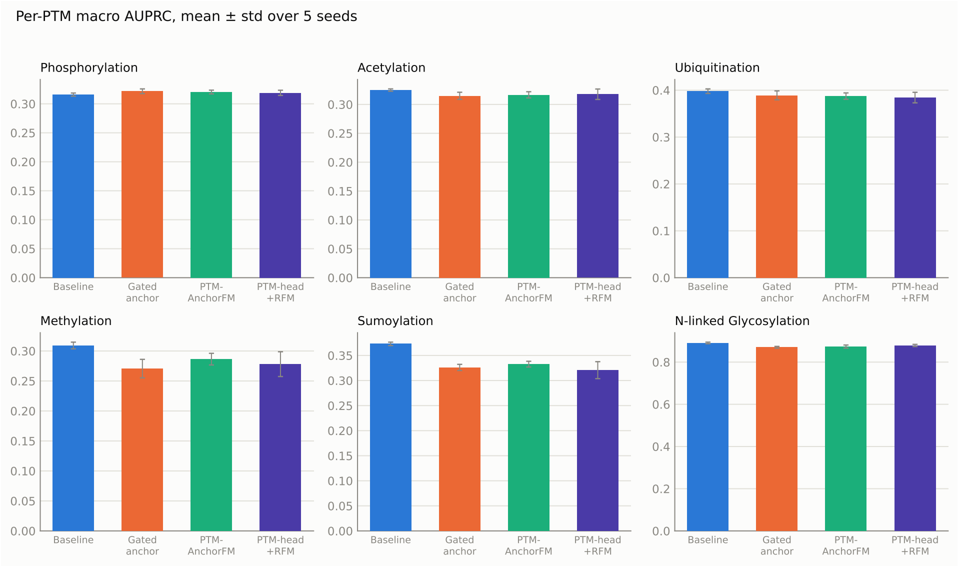


Figure 2: Per-PTM AUPRC with mean and standard deviation over five random seeds.

### D Reproducibility Details

#### D.1 Main Configuration

The main PTM-AnchorFM configuration uses $M = 16$ PTM-positive anchors and $B = 32$ residue-background anchors. Anchors are computed by mini-batch $k$-means after residue-wise standardization of ProtT5 embeddings. Flow pairs are constructed using nearest-neighbor anchor matching, and the trained flow field is evaluated at $t_{\text{eval}} = 0.5$. The flow model is trained once across PTM–residue pairs and used as a frozen feature extractor for final site scoring.

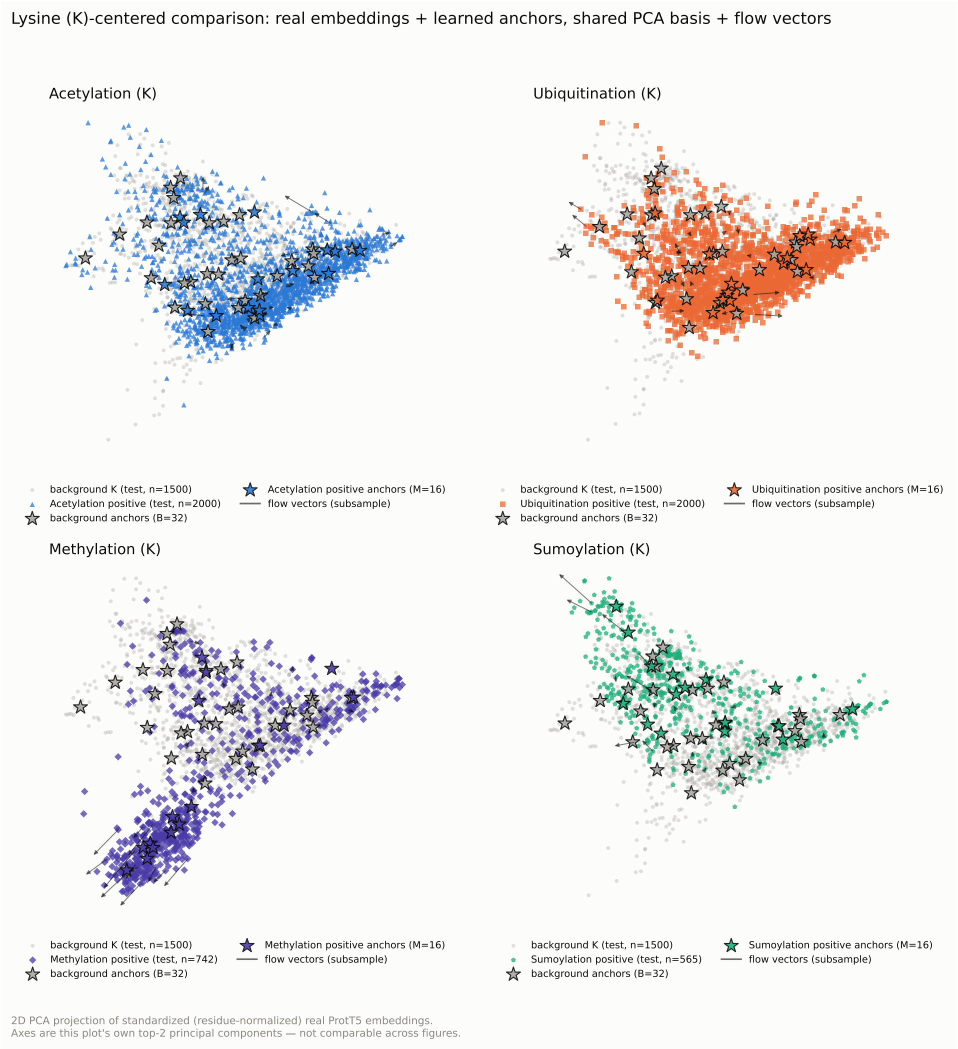


Figure 3: PCA projections of standardized lysine-site embeddings for acetylation, ubiquitination, methylation, and sumoylation. Each panel shows background K residues, PTM-positive K residues, learned anchors, and a subsample of flow vectors.

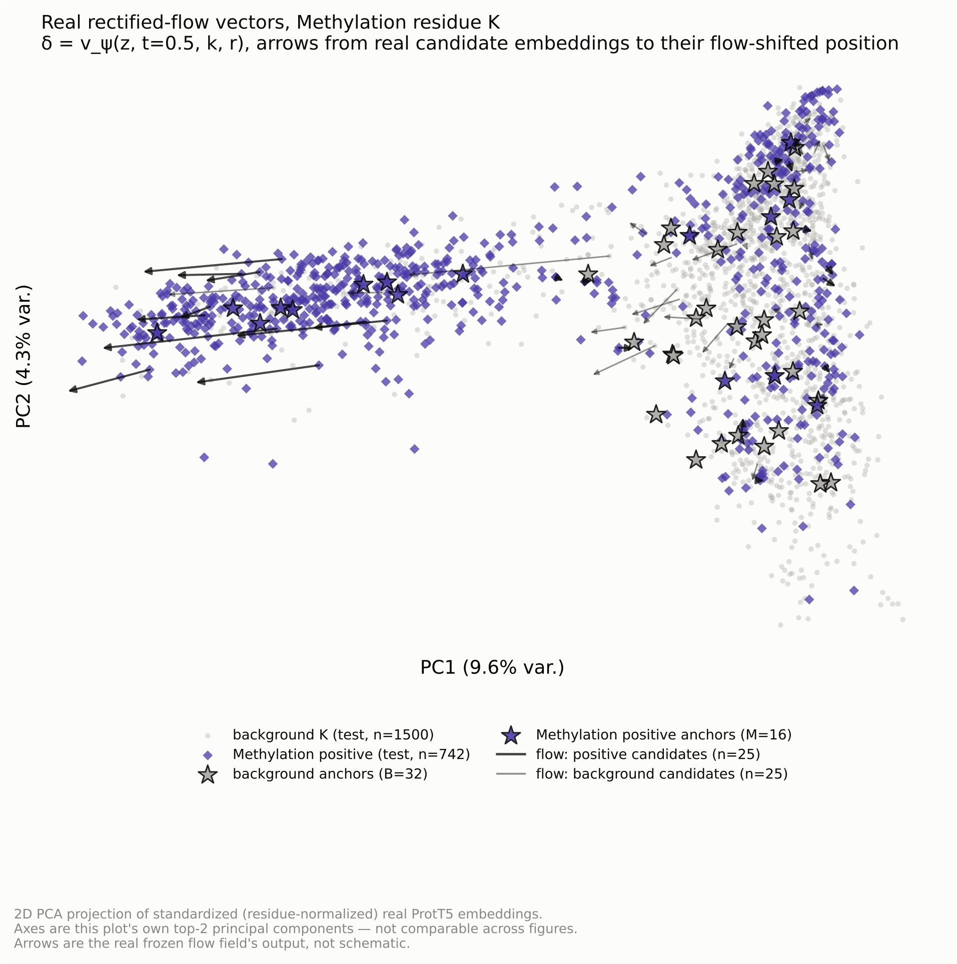


Figure 4: Real rectified-flow vectors for methylation on lysine. Arrows show the frozen flow field output projected into two PCA dimensions.
